# Plasma membrane topography governs the three-dimensional dynamic localization of IgM B cell receptor clusters

**DOI:** 10.1101/2022.04.29.489661

**Authors:** Deniz Saltukoglu, Bugra Özdemir, Michael Holtmannspötter, Ralf Reski, Jacob Piehler, Rainer Kurre, Michael Reth

**Author notes:** **to whom correspondence should be addressed:** Michael Reth.

## Abstract

B lymphocytes recognize bacterial or viral antigens via different classes of the B cell antigen receptor (BCR). Protrusive structures termed microvilli cover lymphocyte surfaces and are thought to perform sensory functions in screening antigen-bearing surfaces. Here, we have studied the cell surface features of Ramos B cells and the spatiotemporal organization of the IgM-BCR using lattice light sheet microscopy in combination with tailored custom-built 4D image analysis. Ramos B cell surfaces were found to form dynamic networks of elevated ridges bridging individual microvilli. A proportion of membrane-localized IgM-BCR was found in clusters, which were associated with the ridges and the microvilli. The dynamic ridge network organization and the IgM-BCR cluster mobility were linked and both were controlled by Arp2/3 complex activity. Our results suggest that topographical features of the cell surface govern the distribution and dynamic localization of IgM-BCR clusters to facilitate antigen screening.

## Introduction

Clonal selection of B lymphocytes is initiated when they recognize cognate antigens via their B cell antigen receptor (BCR) within secondary lymphoid organs [1–4]. In these organs, B cells actively survey specialized antigen-bearing cells [5–13]. Antigen surveillance and sensing are thought to be facilitated by actin-based lymphocyte protrusions called microvilli (MV) that cover the cell surface [7,14–16]. Genetic studies show that components of the actin cytoskeleton play a major role in both the regulation and activation of different BCR classes and that mutations in cytoskeletal genes can cause auto-immune diseases and/or immunodeficiency [17–22]. The IgM- and IgD-class BCRs on naïve B cells form separate clusters on the B cell surface [23]. IgM- and IgD-BCR harbor different transmembrane proteins in their proximity which they functionally connect to, suggesting that the B cell plasma membrane is compartmentalized [24,25], similar to T cells [26,27]. Cell surface topography in conjunction with the actin cytoskeleton has the potential to contribute to the compartmentalization of cell surface molecules. Topography-dependent distribution of immune-related cell surface proteins is a topic that currently arouses interest and has implications for receptor silencing and immune cell surveillance strategies [7,14,28,29]. Live cell total internal reflection (TIRF) microscopy studies suggest that cell surface dynamics of the IgM-class BCR is linked to the underlying actin cytoskeleton [17,30–33]. However, 3D topography-dependent localization of IgM-BCR is difficult to address with TIRF microscopy because imaging is limited to a single plane and the requirement to spread B cells on substrates leads to the flattening of protrusive structures such as the MV. Variations of the TIRF technique such as variable angle-TIRF microscopy performed on fixed T cells demonstrated that the T cell receptor (TCR) and CCR7 accumulate at the tips of MV (MT) [34,35]. An improved substrate coating to keep T cell surface protrusions intact in living cells and application of three-dimensional (3D) single molecule localization microscopy (SMLM) at the cell-substrate interface showed that membrane protrusions segregate CD4 from the transmembrane phosphatase CD45 [28]. With expansion microscopy, it was demonstrated that CD45 is excluded from MT in T and B cells [36].

To understand IgM-BCR plasma membrane localization in the context of dynamic 3D surface topography, we here performed dual color lattice light sheet microscopy (LLSM) of IgM-BCR and a plasma membrane marker on cultured Burkitt’s lymphoma Ramos B cells, on which the function and spatial distribution of many molecules such as CD19, CD20, CXCR4 resemble normal peripheral B cells [24,25,37]. LLSM with its ultra-thin light-sheets allows for high-speed 3D imaging with lowest phototoxicity and photobleaching [38]. It provides almost aberration-free, diffraction-limited resolution with highest sensitivity and unmatched temporal resolution. These features make LLSM an effective tool to study 3D surface dynamics. Quantification of cell surface intensity distributions in light sheet microscopy has previously been performed for PIP2, K-ras and septins [39,40]. In this work, we developed dedicated image analysis methods to correlate receptor localization with cell surface topography. We showed that IgM-BCR forms clusters on the unperturbed 3D surface. We uncovered that the B cell surface is patterned by an elevated ridge network on which MV emerge. IgM-BCR clusters localize to these protrusive structures and couple with their dynamics via the Arp2/3 complex. Our findings imply an active role of surface topography in functional BCR localization to facilitate antigen surveillance.

## Results

### The IgM-BCR is clustered on the Ramos B cell surface

For cell surface-selective photostable fluorescence labeling, we fused 4-hydroxy-3-iodo-5-nitrophenylacetyl (NIP)-specific IgM-BCR to an N-terminal SNAP-tag (SNAP-IgM-BCR). We retrovirally expressed SNAP-IgM-BCR in heavy and light chain gene-knockout (H/L-KO) Ramos B cells [41] along with EGFP-CaaX to visualize the plasma membrane (Suppl. Fig. 1a, b). As control, we used an N-terminally tagged CD40 (SNAP-CD40). We confirmed SNAP-IgM-BCR’s uncompromised signaling competence by activation with antigen (Suppl. Fig. 1c, d). After cell surface labeling of the SNAP-tag, Matrigel-embedded Ramos B cells were imaged by LLSM in two channels (EGFP, DY549P1) with a speed of 2.5 s per volume. To limit the analysis of IgM-BCR and CD40 molecules to the plasma membrane, we created cell surface masks by segmenting the EGFP-CaaX volumes (Fig. 1a, Suppl. Fig. 1e). Segmentation was performed via a modification of nnU-Net [42] using self-generated ground truth. To faithfully extract the surface topography and MV protrusions, the algorithm was developed to be sensitive to fine details (Vid. 1).

**Figure 1:**
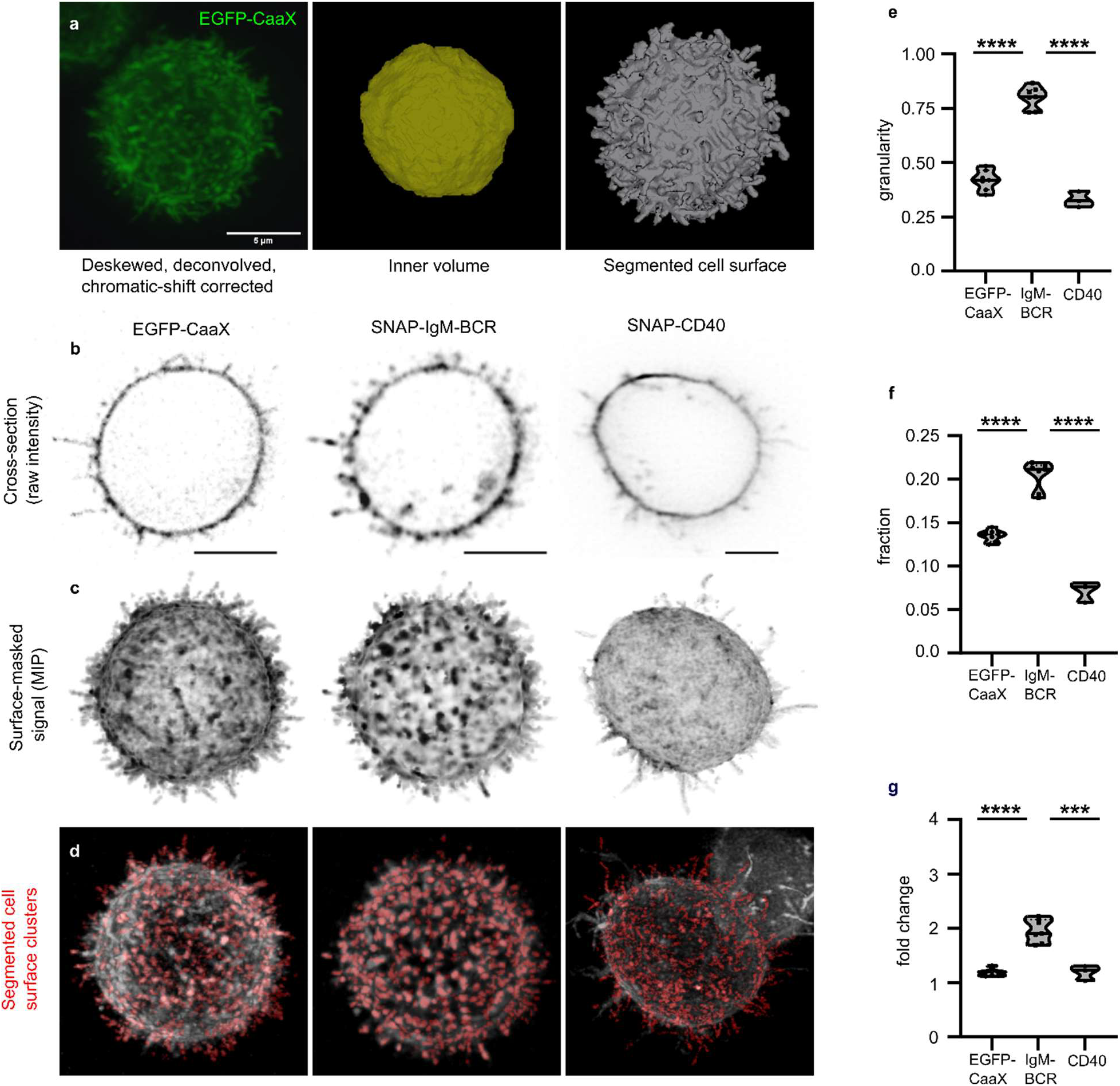
Distribution of SNAP-IgM-BCR and SNAP-CD40 on the B cell surface. **(a)** 3D image of an EGFP-CaaX-expressing Ramos B cell embedded in Matrigel (left) and segmentation of the inner dome (center) using our own script based on Watershed segmentation and segmentation of the cell surface (right) using an nnU-Net model that we trained based on our own-generated ground truth. **(b)** Cross-sectional raw intensity images of SNAP-IgM-BCR/EGFP-CaaX and SNAP-CD40 expressing Ramos B cell. **(c)** Maximum intensity projection of surface-masked image stacks of the cells in **c**. **(d)** Cluster masks generated by custom clustering algorithm based on local contrast. **(e)** Cluster granularity score. **(f)** Total voxel intensity in segmented clusters (Σ _ci_) in comparison to the total voxel intensity within the cell surface mask (Σ _ti_). **(g)** Enrichment of intensity within the clusters. Average voxel intensity in segmented clusters (μ_ci_) in comparison to the average voxel intensity of the non-clustered (“diffuse”) signal within the cell surface mask (μ_di_). n = 7 for EGFP-CaaX and IgM-BCR, n = 3 for CD40. Two-tailed, unpaired t-test was used in **e**, **f**, **g**. p < 0.0001 **** Scale bars = 5 μm.

Visual inspection of the EGFP-CaaX, SNAP-IgM-BCR and SNAP-CD40 signals in cross-sectional slices and surface-masked 3D volumes showed that these proteins are differentially distributed on the plasma membrane (Fig. 1b, c, Vid. 2). Compared to EGFP-CaaX and SNAP-CD40, SNAP-IgM-BCRs accumulated into dot-like, high-contrast structures. To quantify this effect, we developed a cluster segmentation method based on local intensity contrast in 3D volumes. This method extracted distinct clusters of IgM-BCR (Fig. 1d, e, Vid. 3). Along with the cluster segmentation, we proposed a metric (namely, “granularity”) that roughly quantifies the relative accumulation of the signal in clusters in ratio to the more homogeneously distributed signal. After applying our cluster segmentation method to different classes of images, significantly higher granularity scores were obtained for IgM-BCR compared to EGFP-CaaX and SNAP-CD40 (Fig. 1d, e, Vid. 3). In addition to granularity, we also quantified other features to understand whether the IgM signal is enriched within the segmented clusters. We found that the total signal within the segmented IgM-BCR clusters amounted to 22% of the surface-masked total receptor signal with a 2.1-fold increase in average intensity (Fig. 1f, g). This value was lower for EGFP-CaaX (14%) or CD40 (8%) with much less enrichment (1.2-fold) as compared to IgM-BCR (Fig. 1f, g). This quantification highlights that a substantial fraction of the IgM-BCR complexes on the plasma membrane of resting-state B cells reside in clusters.

### 3D plasma membrane topography of Ramos B cells

To understand the cell surface topography of B cells in more detail, we represented the surface mask as a triangular mesh and computed the micron-scale shapes using the shape index algorithm [43]. The color-coded index shows that the B cell surface can be described by shallow invaginating cups (in yellow, labelled as “c”) surrounded by saddles and elevated ridges (in blue, labelled as “r”) in a repetitive way (Fig. 2a, b). Thus, the whole Ramos B cell surface was classified into a pattern of positively- and negatively-curved micro-topographies. This classification revealed a continuous network of elevated ridges on the B cell surface, from which MV protrusions emerge (in purple, labelled as “m”) (Fig. 2a, b). To analyze the organization of the ridge structure in more detail, we used a Hessian-based line filter to mask the curvilinear features on the raw intensity images of EGFP-CaaX (Suppl. Fig. 1e). When the outcome (excluding the MV) was superimposed onto the EGFP-CaaX image, the extracted lines correlated with the elevated regions of the cell surface (Fig. 2c). These elevated regions displayed higher EGFP-CaaX intensities, possibly because the ridges create opposing membranes that are not resolved due to the diffraction limit of light and/or because they are preferred by the CaaX motif.

**Figure 2:**
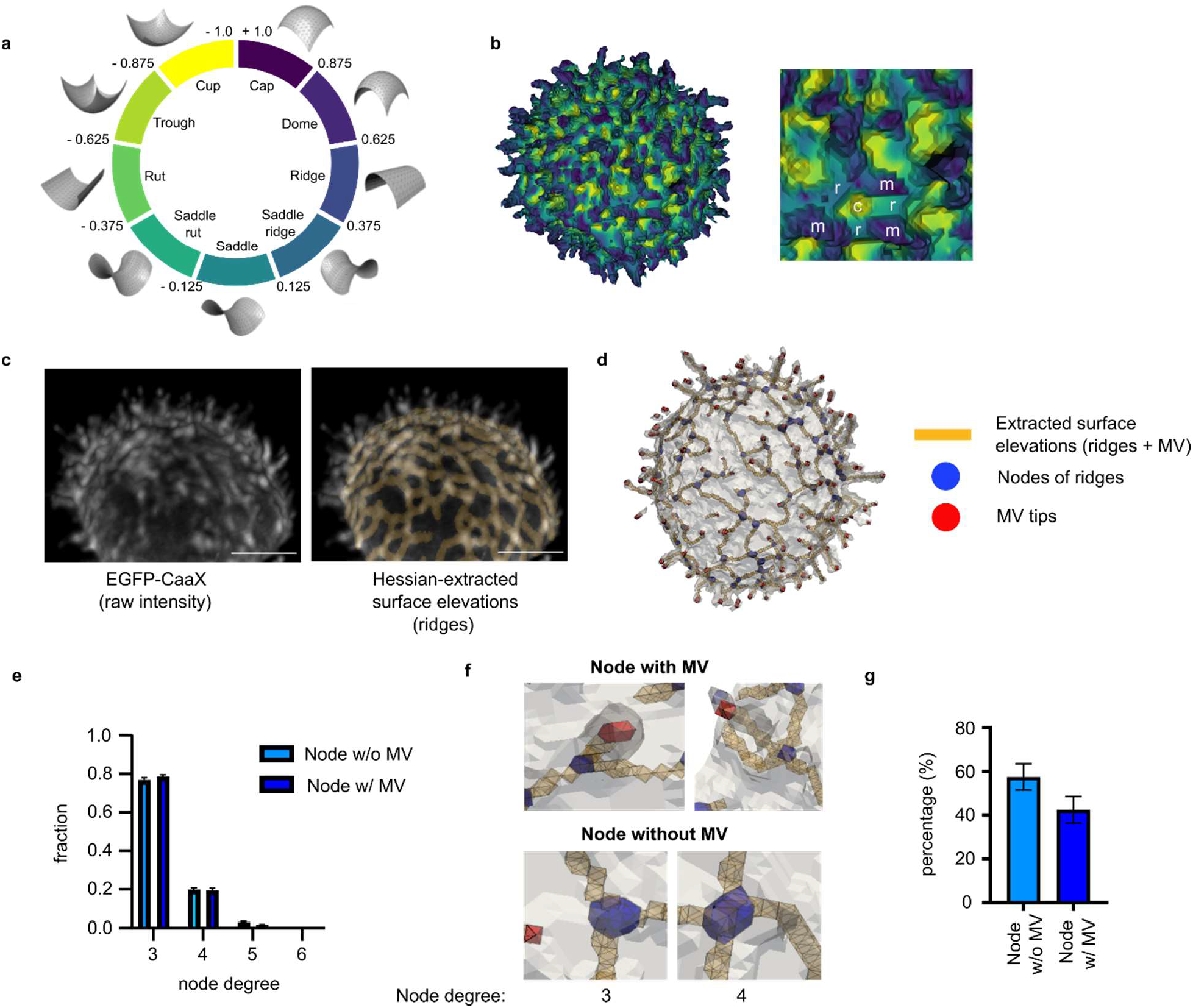
Identification of distinct morphological features on the B cell surface. **(a)** Color code of shapes according to the shape index algorithm. **(b)** Application of the shape index to the triangular surface mesh to highlight surface structures on the B cell. The zoomed area reveals individual MV (m) connecting to each other via positively curved ridges (r) and saddles, surrounding positively curved cups (c) in a patterned way. **(c)** Extraction of the ridge network on the B cell surface. Positively curved ridges were extracted from raw EGFP-CaaX images (left) using a Hessian-based algorithm and superimposed on the raw image (right). Scale bar, 5 μm. **(d)** The skeletonized ridge network including the MV protrusions superimposed on the mesh representation of the cell surface. The ridges, their connecting nodes and the MV tips are indicated by the yellow, blue and red color, respectively. **(e)** Node degree analysis of the skeletonized ridge network. **(f)** Enlarged images of **d** displaying exemplar 3- and 4-degree nodes with or without MV association. **(g)** Percentage of the nodes without or without MV association.

The skeletonized form of the line mask (including the MV) superimposed on the surface mesh formed an interwoven network throughout the cell surface (Fig. 2d). Importantly, all MV structures on the cell surface were extracted in connection with the ridges, confirming that MVs are extensions on the elevated ridge network. Network node analysis showed that most of the network nodes with or without a MV connection have a node degree of 3, implying simple branching in their formation (Fig. 2e, f). In a smaller fraction (20%) of the nodes, the node degree was 4. Nodes with MV connection comprised 40% of all the extracted nodes on the skeleton (Fig. 2g). In sum, the Hessian line extraction analysis showed that the elevated ridges and MV form an interconnected network that is topographically distinct from the rest of the B cell surface.

### IgM-BCR clusters are enriched on the ridges and microvilli

After detailing the cell surface topography of Ramos B cells, we investigated whether IgM-BCR clusters are enriched at a particular surface structure. To do so, we compartmentalized the cell surface into distinct topographical features, namely, shallow invaginations (SI), ridges (R), nodes (N), MV shafts (MS) and MV tips (MT) (Fig. 3a, Suppl. Fig. 1e). N corresponded to the sites where R and MV connect (MV roots) or to the sites where R laterally branch. Among the surface features, MV occupied the most volume, followed by SI and (R + N) (Suppl. Fig. 2a).

**Figure 3:**
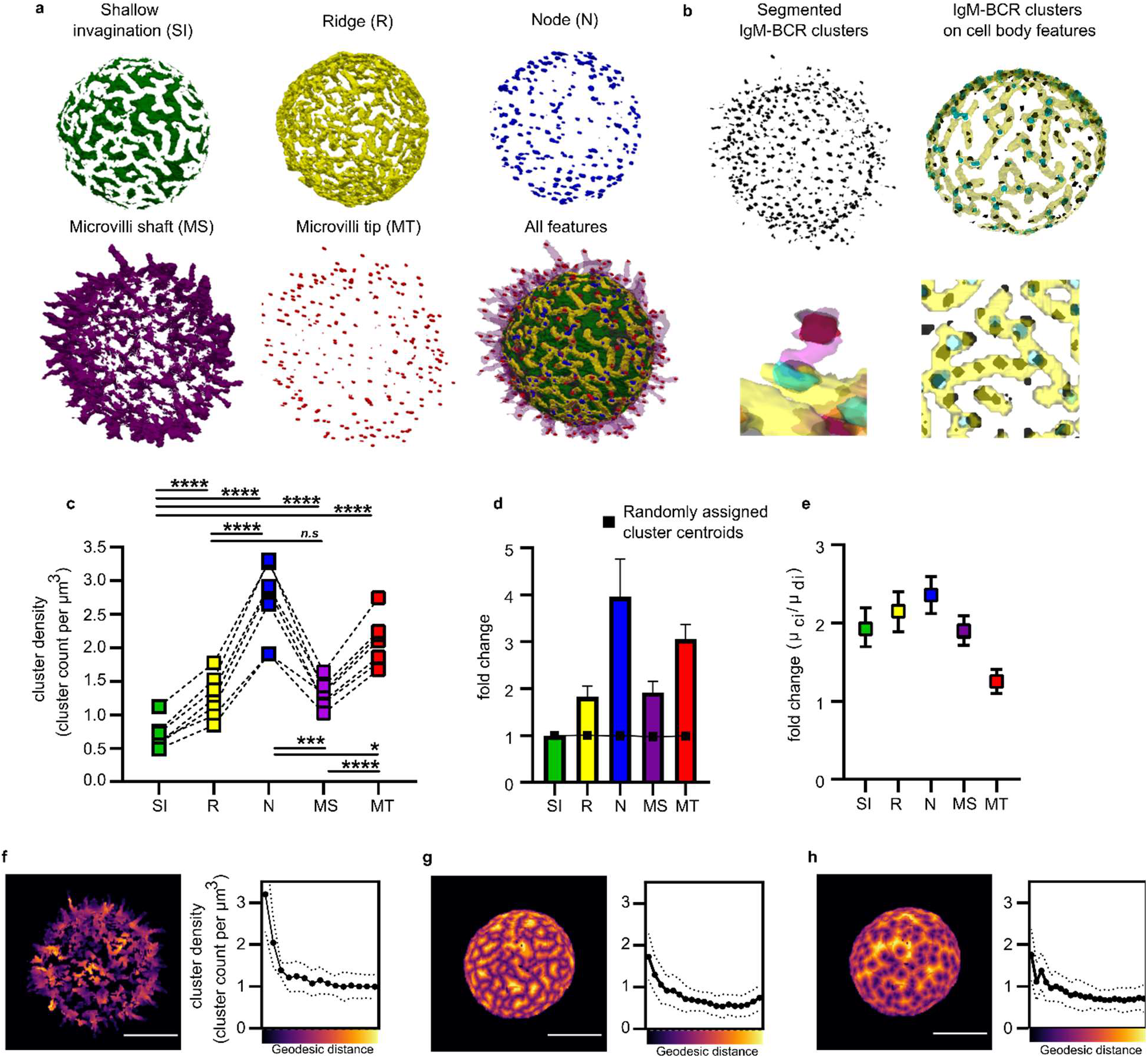
IgM-BCR clusters are enriched on the ridges and microvilli. **(a)** Topographical features of the B cell surface. The cell body is separated from the MV protrusions using mathematical morphology. On the cell body, the ridge (R) network (yellow) is computed and its nodes (N) (blue) are annotated. The remaining surface area is termed shallow invaginations (SI, dark green). Individual MV are split into their shaft (MS, purple) and tip (MT, red) and all features are combined in one image. **(b)** Segmented IgM-BCR clusters are superimposed onto the extracted surface features. Cluster-surface feature overlay reveals the accumulation of IgM-BCR clusters on the ridge network. The zoomed-in MV displays a cluster density at its root (corresponding to a N) and tip. **(c)** Cluster density at surface features. IgM-BCR cluster density within different topographical features quantified as cluster count per surface feature volume. 7 cells (30-time frames/cell) were analyzed and subjected to two-tailed, paired student t-test. p < 0.5 *, p < 0.01 **, p < 0.001 ***. **(d)** Normalized cluster density. Comparison of mean cluster densities computed in **c** as normalized to the value of SI. The black squares indicate the SI-normalized cluster density of randomly assigned cluster centroids on the cell surface. **(e**) IgM-BCR intensity enrichment within the segmented clusters at different surface features, calculated for the cells quantified in **c**. Intensity enrichment was measured by dividing the average voxel intensity in the IgM-BCR clusters (μci) by the average voxel intensity of the non-clustered IgM-BCR residing within the surface feature mask (μdi). **(f, g, h)** Continuous analysis of IgM-BCR cluster density using geodesic distance maps initiated from **(f)** MT towards MV roots, **(g)** centerline of R towards cell body surface, and **(h)** N towards cell body surface.

To quantify cluster density at each surface feature (cluster count / feature volume), we represented each segmented IgM-BCR cluster by the coordinates of its centroid. Simulated random coordinates on the surface mask were distributed evenly across the five different surface features, showing that feature volume does not bias distribution (Fig. 3d). In contrast, IgM-BCR clusters were least dense within the SI and 1.8- and 4-fold enriched on the R and N, respectively (Fig. 3b, c, d). Within the MV, cluster density was 1.5-fold enriched at the MT in comparison to MS (Fig. 3b, c, d).

To account for the extent of IgM-BCR accumulation in clusters at different surface features, we calculated the ratio of the mean IgM-BCR intensity within clusters (μ_ci_) to the mean non-clustered (“diffuse”) IgM-BCR intensity from the whole surface (μ_di_). This ratio was highest at N with 2.5-fold increase and lowest at MT with a 1.2-fold increase (Fig. 3e). Since our clustering algorithm detects local contrast, it can efficiently identify clusters within a high (R) and low (MV) diffuse intensity background (Suppl. Fig. 2b, c, d).

In addition to this discrete analysis, we assessed the cluster distribution in a continuous manner with geodesic distance maps. We initiated these maps either from the R centerline, the N or the MT and moved out, sampling equal volumes (Fig. 3f, g, h). Along the length of the MV geodesic distance map, IgM-BCR cluster density was highest at MT (Fig. 3f). The IgM-BCR cluster density at the R and N geodesic distance maps also displayed a sharp decrease moving away from the initiation sites (Fig. 3g, h). As a control, the decrease in CD40 cluster density was not as steep when moving away from R centerline or N (Suppl. Fig. 2f, g). However, CD40 density was also highest at the MT within the MV (Suppl. Fig. 2e). These analyses coherently substantiate that IgM-BCR clustering is localized in a patterned fashion and enriched on certain topographical features.

### IgM-BCR mobility and R network dynamics are linked by Arp2/3 complex activity

Since the R network predominantly has a node-degree of 3, it is possible that branched actin polymerization mediated by the Arp2/3 complex is involved in its formation or dynamics. In cells treated with 100 μM of the Arp2/3 complex inhibitor CK-666 for 30 min, phalloidin staining did not detect significant decrease in polymerized cortical actin (Suppl. Fig. 3a). In cells treated with CK-666 for 15 – 45 min, the R network could still be extracted and had a similar node-degree distribution as in untreated B cells (Suppl. Fig. 3b). For visual inspection of the dynamics, we produced a sum projection of the skeletonized R network and MV along the time dimension in 15 consecutive frames, covering a window of 37.5 s. For a quantitative analysis of the dynamics of surface features and clusters, we propose the metric “staticity”. Staticity is inversely correlated with the dynamics of the feature mask being analyzed. For the surface feature images, the staticity was calculated based on similarity scores using the “intersection over union” method [44] for consecutive pairs of time frames and was separately calculated for the R and MV. For cluster centroids the staticity was calculated from a staticity heatmap which was derived by convolving the datasets of cluster centroids with a 4D Gaussian kernel.

Untreated cells yielded blurred sum projection images of R and MV with low staticity scores, highlighting their dynamic nature (Fig. 4a, d). In stark contrast, CK-666-treated cells displayed a frozen R network with high staticity scores. Different from what is reported for Jurkat T cells, the cumulative surface coverage of dynamic cell projections seemed to depend more on R dynamics rather than lateral MV movements in Ramos B cells (Suppl. Fig. 3c) [7]. CK-666 effectively abolished the surface coverage mediated by the R network.

**Figure 4:**
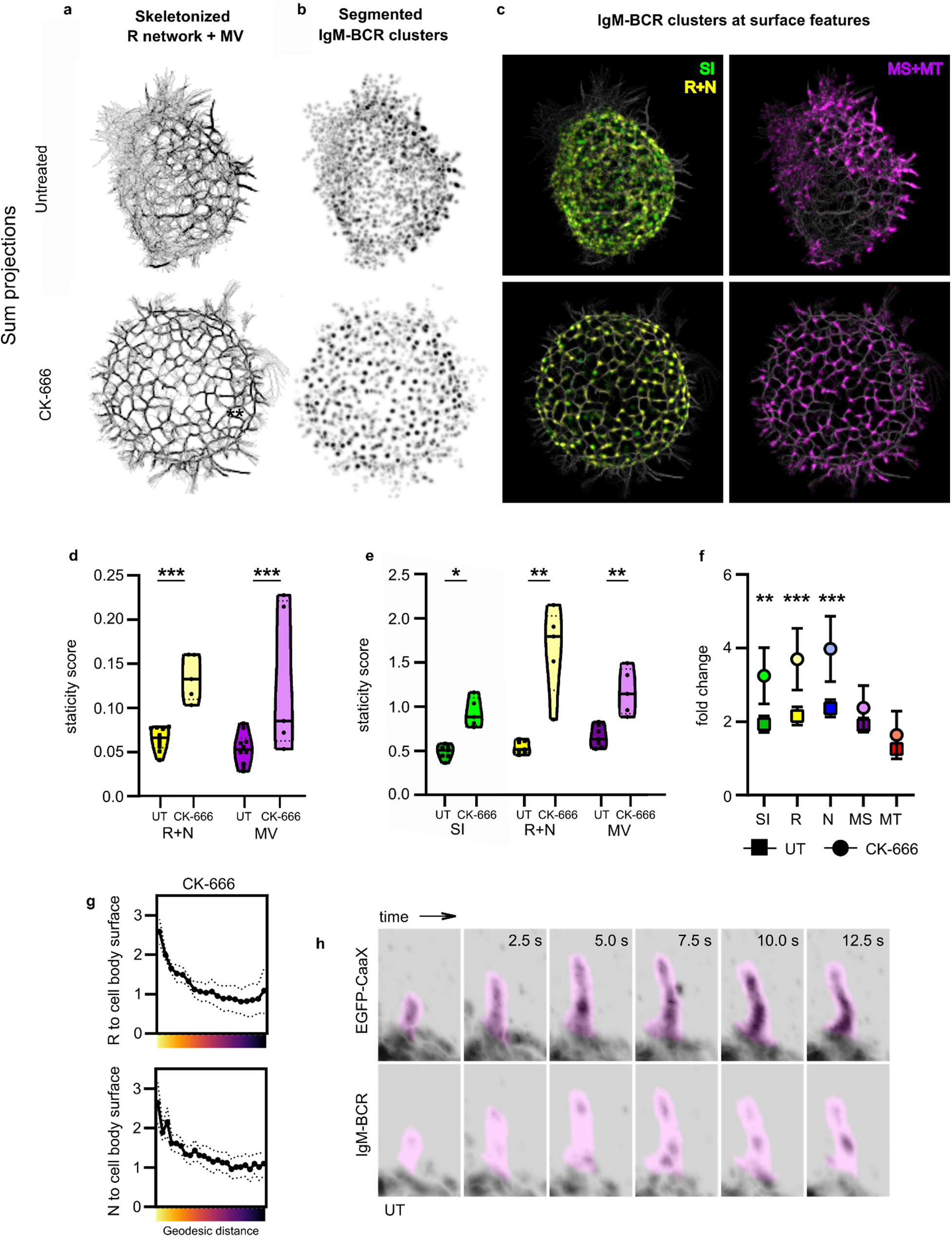
IgM-BCR cluster dynamics are linked with R network dynamics in an Arp2/3 complex-dependent manner. **(a, b)** Sum projections of the skeletonized R network and MV **(a)** and segmented IgM-BCR clusters **(b)** in an untreated and 100 μM CK-666-treated cell for 15 consecutive time frames amounting to 37.5 s. **(c)** Color coding of the segmented IgM-BCR clusters in **(b)** according to their assigned surface feature. Yellow clusters correspond to SI, green to R + N, purple to MV and MT. **(d, e)** Staticity scores for different surface features (R+N, MV) **(d)** and for segmented IgM-BCR clusters assigned to different surface features **(e)** in untreated and 15 – 30 min 100 μM CK-666-treated cells. Staticity score is determined by calculating the proportion of the objects that can be linked over 3-time frames. **(f)** IgM-BCR intensity enrichment within the segmented clusters at different surface features. Intensity enrichment was measured by dividing the average voxel intensity in the IgM-BCR clusters (μci) by the average voxel intensity of the non-clustered (“diffuse”) IgM-BCR residing within the surface feature mask (μdi). **(g)** Continuous analysis of IgM-BCR cluster density in 15-30 min 100 μM CK-666-treated cells using geodesic distance maps initiating from distinct surface features (centerline of R, N and MT) (to compare to untreated in Fig. 3f). **(h)** Time-course images of EGFP-CaaX and IgM-BCR intensity in an elongating MV (purple, MV mask). n = 7 for untreated cells and n = 4 for CK-666 treated cells. Two-tailed, unpaired student t-test was used in **d, e, f**. p < 0.5 *, p < 0.01 **, p < 0.001.

Interestingly, CK-666 treatment also strongly affected clustering of IgM-BCR on the topographic landscape of the B cell. In comparison to untreated cells, Arp2/3 complex-inhibited cells displayed a higher IgM-BCR cluster density at R and N (Fig. 4g, 3f). These clusters were brighter (Fig. 4f) and arrested in mobility (Fig. 4b, c, e, Vid. 4). In some frames, arrested clusters were assigned to the roots of MV (Fig. 4c). It is likely that these clusters occupy the highly curved joints at the R and MV intersection, suggesting that Arp2/3 complex function is required for IgM-BCR cluster mobility at this topographical region. During MV elongation, IgM-BCR entered the MV in a clustered form (Fig. 4h, Vid. 5). IgM-BCR clusters dissipated and reformed, however, in a way that seemed to produce a forward movement towards the MT. In Arp2/3 complex-inhibited cells MV neither underwent elongation, nor de-novo formation (Vid. 4). Taken together, these data suggest that Arp2/3 complex-dependent cytoskeletal dynamics may promote the outward translocation of IgM-BCR clusters from the N pool to MV structures and link IgM-BCR cluster mobility to R and MV dynamics.

### Antigen-induced micro-clusters associate with the ridge network

To find out how antigen stimulation affects IgM-BCR distribution on cell surface features, we used 50 ng/ml of 15mer NIP coupled to BSA (NIP-15-BSA) to stimulate the cells for 5 – 30 min. The binding of the cognate antigen led to the formation of the typical bright antigen-induced IgM-BCR micro-clusters (Ag-MC) (Fig. 5a). Using high-stringency cluster extraction (with local intensity fold change 3x or higher), we found that these Ag-MC concentrated on the cell body (SI, R, N) but not on MT and MS (Fig. 5a, b). In terms of Ag-MC and R network dynamics, there were two profiles. In one profile, Ag-MC were immobilized on a frozen R network (Lower panel of Fig. 5c, d, e) with higher cluster and R network staticity scores (Fig 5g, h). Interestingly, these cells displayed local R mobility at areas that lacked an Ag-MC (Fig. 5f). In the second profile, Ag-MC underwent displacement on non-stabilized R (Upper panel of Fig. 5c, d, e, g, h, i). Therefore, surface topography provides a platform for the anchoring and directional movement of Ag-MC.

**Figure 5:**
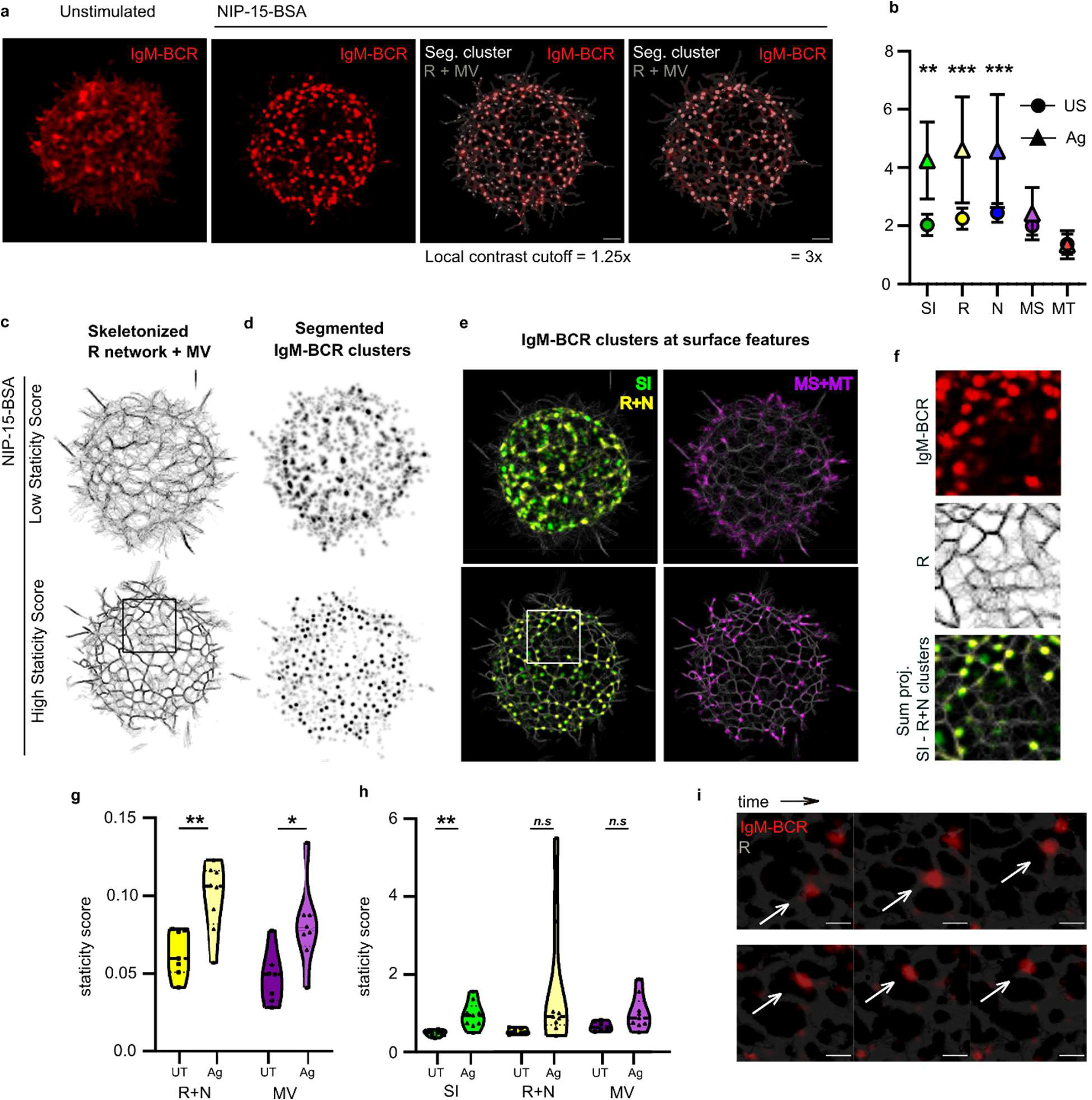
Antigen-induced micro-clusters (Ag-MC) localize to and move on the R network. **(a)** IgM-BCR cluster distribution upon Ag stimulation. Ramos B cells in the Matrigel were stimulated with 50 ng/ml NIP-15-BSA Ag for at least 15 minutes. Ag-MC on the B cell surface were segmented and overlaid with the R network (including MV) with a 1.25x and 3.0x local intensity cut-off for cluster detection. Scale bar, 1 μm. **(b)** IgM-BCR intensity enrichment within the segmented clusters at different surface features for unstimulated (US) and Ag-stimulated cells. Intensity enrichment was measured by dividing the average voxel intensity in the IgM-BCR clusters (μ_ci_) by the average voxel intensity of the non-clustered (“diffuse”) IgM-BCR residing within the different surface feature masks (μ_di_). **(c, d)** Sum projections of the skeletonized R network and MV **(c)** and segmented IgM-BCR clusters **(d)** in Ag-stimulated cells with a low and high staticity score (duration: 37.5 s) **(e)** Color coding of the segmented IgM-BCR clusters in **(d)** according to their assigned surface feature. Green clusters correspond to SI, yellow to R + N, purple to MV and MT. **(f)** Close-up of the encircled region in **e** displaying IgM-BCR localization, sum projections of the R network and segmented clusters. R network lacking Ag-MC are not stabilized. **(g, h)** Staticity scores for different surface features (R+N, MV) **(g)** and for segmented IgM-BCR clusters assigned to different surface features **(h)** in unstimulated and Ag-stimulated cells. Staticity score is determined by calculating the proportion of the objects that can be linked over 3-time frames. **(i)** Time course images of an Ag-MC undergoing mobility on R. n = 7 for ustimulated cells and n = 8 for Ag-stimulated cells. Two-tailed, unpaired student t-test was used in **b, g, h**. p < 0.5 *, p < 0.01 **, p < 0.001.

### IgM-BCR association with elevated topography is upstream of the actin cytoskeleton

Next, we sought to find out how R and MV on the B cell surface are affected by the loss of the cortical actin cytoskeleton. The treatment of Ramos B cells with 2 μM LatA smoothened a large part of the B cell surface (Fig. 6a, b, h). Some R- and spot-like elevations were still present on these cells that also displayed large polarized protrusive regions. Interestingly, the R- and spot-like membrane elevations were mobile and could merge on the plasma membrane (Fig. 6b, Vid. 6). Of note, phalloidin staining demonstrated that 2 μM LatA treatment fully depolymerized the cortical actin cytoskeleton (Fig. 6c). Thus, disorganized proto-R structures could form on B cells lacking an intact cortical actin cytoskeleton.

**Figure 6:**
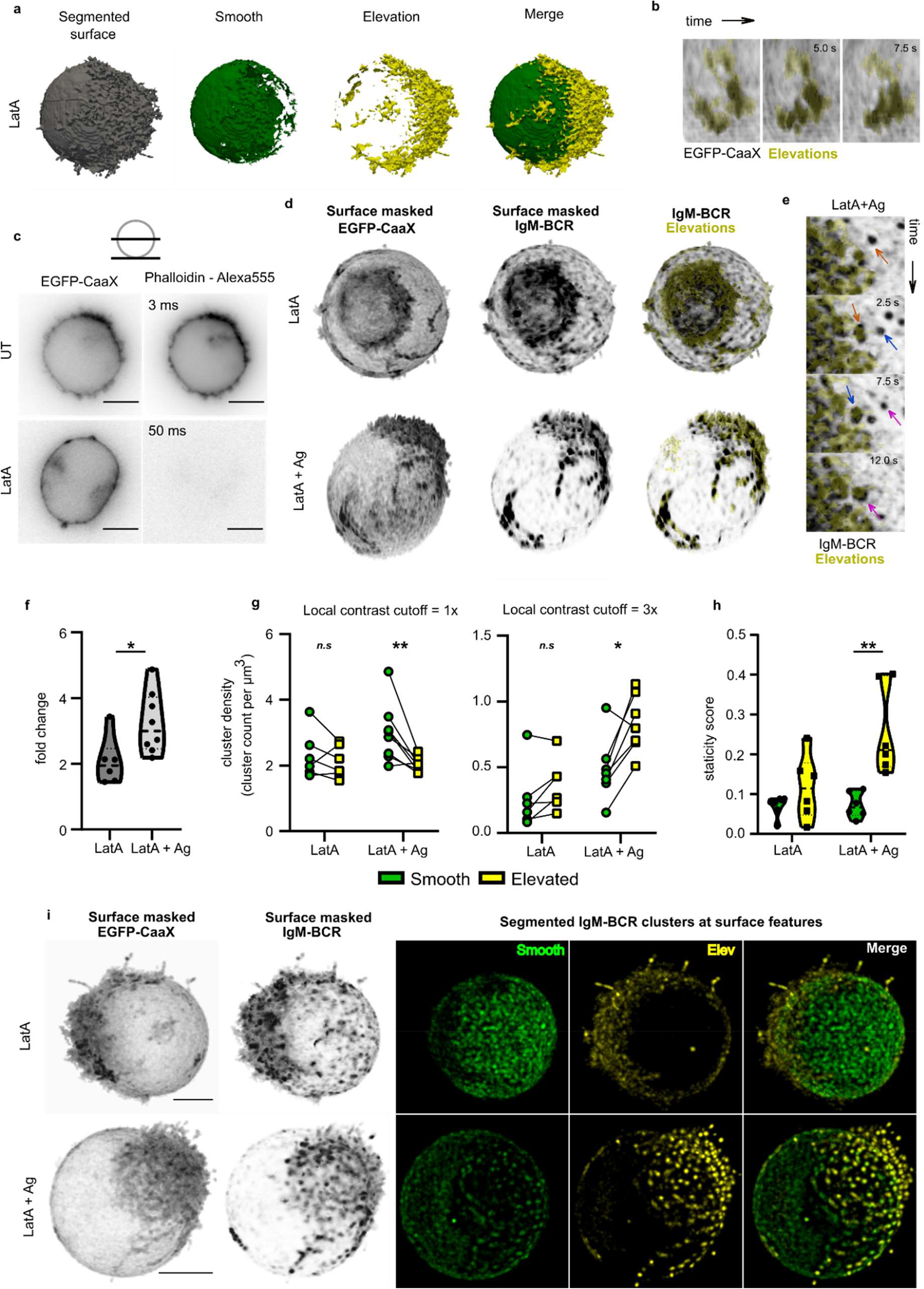
LatA-induced surface elevations anchor Ag-MC. **(a)** Topographical features of the Ramos B cell surface in a cell treated with 2 μM LatA for 15 – 30 min. LatA-treated cells display disorganized and polarized elevations. The cell is morphologically separated into smooth (green) and elevated (yellow) regions. **(b)** Time frames from a LatA-treated cell in which elevations undergo merging. **(c)** Phalloidin-Alexa555 staining of fixed and permeabilized Ramos B cells expressing EGFP-CaaX and IgM-BCR with or without 30 min 2 μM LatA treatment. **(d)** Surface-masked EGFP-CaaX and IgM-BCR signal and overlay of segmented elevations (yellow) with the surface-masked IgM-BCR signal in cells treated with LatA and LatA+Ag (50 ng/ml NIP-15-BSA). **(e)** Time frames from a LatA+Ag cell showing directed movement of Ag-MC towards the elevations (yellow). Each Ag-MC shown with a different colored arrow. **(f)** IgM-BCR intensity enrichment within all the segmented clusters in LatA and Lat+Ag cells. **(g)** Cluster density at surface features. Quantification of the IgM-BCR cluster density in terms of cluster count per surface feature volume for all segmented IgM-BCR clusters (left panel, local contrast cutoff = 1x) and for Ag-MC (right panel, local contrast cutoff = 3x). **(h)** Staticity scores for segmented IgM-BCR clusters on smooth and elevated regions in LatA and LatA+Ag cells. **(i)** Surface-masked EGFP-CaaX and IgM-BCR signal for a LatA and LatA+Ag cell and the sum projections of their segmented IgM-BCR clusters (duration: 37.5 s). IgM-BCR clusters are color-coded according to their assigned surface region. Scale bar, 5 μm. n = 7 for LatA cells and n = 7 for LatA+Ag cells. Two-tailed, paired student t-test was used in **g** and two-tailed, unpaired student t-test was used in **f**, **h**. p < 0.5 *, p < 0.01 **.

With LatA treatment, IgM-BCR clustering was not lost. In smooth regions, as the topographical unevenness of the surface was lost, the differences in the intensity distribution of EGFP-CaaX and IgM-BCR became more evident (Fig. 6d, i, Vid. 7 - upper panel). Segmented IgM-BCR clusters in LatA-treated cells did not display a preference for smooth or the elevated regions (Fig. 6g). However, after antigen stimulation of LatA-treated B cells, Ag-MC were distinctively enriched on the elevated regions (Fig. 6d, g, Vid. 7 - lower panel). This enrichment seemed to result from directional mobility of Ag-MC towards the elevated regions (Fig. 6e, Vid. 8), similar to what is observed in the formation of the immunological synapse. Ag-MC became immobilized on the elevated regions, displaying high staticity scores and bright cluster centroid sum projection images (Fig. 6h, i). The smoothening of the cell surface did not disseminate IgM-BCR clusters but resulted in rapidly moving clusters that lacked the coordination that the R network provided (Vid. 7 - upper panel). However, interestingly, sum projection of mobile clusters displayed a pattern formed by linear trajectories and cluster-absent regions (Fig. 6i). In conclusion, the basic patterning for IgM-BCR cluster localization, directional movement and anchoring of Ag-MC are regulated by mechanisms upstream of the actin cytoskeleton.

## Discussion

Analyzing the Ramos B cell surface in 4D, apart from the well-documented lymphocyte MV [14,16], we observed an elevated, continuous and dynamic surface structure that interconnects neighboring MV, which we termed the ridge network. A similar structure, named micro-ridges, has been found on blood-purified lymphocytes by scanning electron microscopy [45]. Cumulative surface coverage of mobile MV has been proposed as a mechanism in Jurkat T cells to screen opposing membranes for antigen [7]. In Ramos B cells, R covers more surface area in a given time span compared to MV. R dynamics and IgM-BCR cluster distribution on topographical features seem to be interconnected via Arp2/3 complex activity. Inhibition of the Arp2/3 complex blocks R restructuring, halts MV elongation and leads to arrested, larger IgM-BCR clusters, most notably at MV roots (N). In untreated cells, IgM-BCR enters into elongating MV in clusters and makes a forward movement towards MT. We propose that Arp2/3 complex activity is necessary for the coordinated localization of IgM-BCR on MV and R functions as a receptor reservoir. In lymphocytes, where the cellular polarity is subjected to constant reorganization, the association of IgM-BCR clusters with a dynamic structure that controls MV positioning may solve the problem of IgM-BCR availability within cellular protrusions to facilitate antigen sensing. We further speculate that MV elongation may be regulated by external signals warning B cells of an infection or by soluble factors from specialized antigen-bearing cells. This is in line with the finding that TLR4 signaling increases IgM-BCR mobility and promotes cofilin activity, a molecule involved in MV elongation [46,47]. Movement of IgM-BCR clusters from MV roots towards the MT could further function to reduce the threshold for BCR activation, as the transmembrane phosphatase CD45 that negatively regulates BCR activation has been found to be excluded from MT [36].

Our LatA experiments have shown that neither the actin cytoskeleton, nor the topographical features are responsible for the clustering of IgM-BCR. However, these factors are involved in the coordinated movement of the receptor clusters. In cells devoid of the actin cytoskeleton, IgM-BCR cluster mobility is fast and erratic. Nonetheless, their trajectory on the plasma membrane still seems to follow a pattern. Such molecular patterns are observed in the positive and negative feedback loops operating on Rho GTPases even in the absence of the cortical actin cytoskeleton [48,49]. It is conceivable that a similar mechanism creates the backbone of the R network by recruiting curvature-inducing BAR domains [50–53] and the actin polymerization machinery [54–57], which then coordinates the functional localization and transport of the BCR on the 3D surface.

We propose that the germinal center-derived Ramos B cell is a good model system to study questions related to MV and surface topography, as it expresses many of the MV-associated molecules common to lymphocytes (Suppl. Table. 1a) [58]. Regulation of lymphocyte surface structures is a fairly unexplored area and many of the molecules discovered in the context of intestinal MV and stereocilia are not expressed by lymphocytes (Suppl. Table. 1b). We think that insight about antigen sensing in connection with the 3D compartmentalization of the B cell surface may open new avenues for better vaccine design.

## Materials and Methods

### Ramos B cell line culture

Ramos B cells were cultured in RPMI-1640 Medium with GlutaMAX™ (Art. No. 61870010, ThermoFisher Scientific) supplemented with 10% fetal calf serum (PAN), 10 U/ml penicillin-streptomycin (Invitrogen) and 10 mM HEPES (Invitrogen). All cell lines were cultured in a 37° incubator with 5% CO_2_.

### Cloning

#### SNAP-IgM-BCR

Geneblocks (IDT) encoding the b1-8 leader sequence and EGFP was amplified with the primer pair CCCTCGTAAAGAATTCATGGGATGGAGCTGTATCATCCTC and GTCGGCGAGCTGACG. Geneblocks encoding the linker with the A(EAAAK)_3_A amino acid sequence and the heavy chain variable region specific for the hapten NIP was amplified with the primers GCGTGCAGCTCGCCGACCACTACCAGCAGAACACCCC and TGAGGAGACTGT GAGAGTGGTGC. Plasmid encoding the mouse μ heavy chain was amplified with the primers TCTCACAGTCTCCTCAGAGAGTCAGTC and GAGGTGGTCTGGATCCTCATTTCACCTTGAACAGGGTGAC. The three fragments were fused using the in-fusion enzyme (Takara) into a pRetroX-TetOne-Puro (Takara) backbone using the EcoRI and BamHI restriction sites. Next, the region encoding the three fused sequences were amplified with GAGGTTAACGAATTCATGGGATGGAGCTGTATCATC and CCAGCCTGCTTCAGCAGGCTGAAGTTAGTAGCTCCGCTTCCTGATTTCACCTTGAACAGGG TGACG. The mouse λ light chain with specificity for NIP was amplified with the primers TGCTGAAGCAGGCTGGAGACGTGGAGGAGAACCCTGGACCTATGGCCTGGATTTCACTTATACTCTCTC and TGGCTGCAGGTCGACCTAGGAACAGTCAGCACGGGA. The two fragments were fused using the in-fusion enzyme into a pM retroviral vector cut with EcoRI (NEB) and SalI (NEB) restriction enzymes. The primers contained the coding sequence for the P2A cleavage site in between the μ heavy and λ light chains. Finally, b1-8 leader-EGFP was switched to a b1-8 leader-SNAP-tag by amplifying the SNAP-tag sequence with GAGGTTAACGAATTCATGGGATGGAGCTGTATCATCCTCTTCTTGGTAGCAACAGCTACAGGTGTCCACTCCATGGACAAAGACTGCGAAATGAAGCG and TCTGCTGGCGGCCGACCCAGCCCAGGCTT and cloning into the plasmid with the in-fusion enzyme using the EcoRI and NotI (NEB) restriction sites before and after the b1-8 leader-EGFP sequence.

#### EGFP-CaaX

EGFP sequence was amplified with the primers CGAGGTTAACGAATTCcATGGTGAGCAAGGGC and attttatcgataagcTTctacataattacacactttgtctttgacttctttttcttcttttta. The amplified fragment was cloned into pM retroviral vector cut with EcoRI and SalI restriction enzymes using the in-fusion enzyme.

#### SNAP-CD40

cDNA encoding CD40 was amplified from the cDNA pool of Ramos B cells with the primer pair CGAGGTTAACGAATTCATGGTTCGTCTGCCTCTGC and GCCGCTTCCGCCTCCCTGTCTCTCCTGC. EGFP was amplified with the primers GGAGGCGGAAGCGGCGTGAGCAAGGGCGAGG and ATTTTATCGATAAGCTTTCACTTGTACAGCTCGTCCATGCC. The two fragments were inserted into a pM retroviral vector cut with EcoRI and HindIII (NEB) with the in-fusion enzyme. Next, CD40 sequence was amplified with GGAGGCGGAAGCGGCGAACCACCCACTGCATGC and TTTTATCGATAAGCTTTCACTGTCTCTCCTGCACTGAGATG. SNAP-tag sequence with an N-terminal b1-8 leader sequence was amplified with CGAGGTTAACGAATTCATGGGATGGA and GCCGCTTCCGCCTCCACCCAGCCCAGGCTG. The two fragments were cloned into a pM retroviral vector cut with EcoRI and HindIII using the in-fusion enzyme.

### Generation of the Ramos B cell lines with retrovirus

H/L chain locus of Ramos B cells was silenced using the CRISPR-Cas9 technology. For a detailed protocol, refer to [41]. The retroviral plasmids encoding SNAP-IgM-BCR, EGFP-CaaX and SNAP-CD40 were transfected (1000 ng) along with the pKat plasmid encoding the viral coat protein (500 ng) into the ecotropic virus generating cell line PlatE using the transfection reagent PolyJet (Cat No. SL100688, SignaGen) in a 6-well plate format. PlatE cells were maintained in the RPMI cell culture medium. The virus containing supernatant was collected for each plasmid after 48 hr. 500,000 H/L-KO Ramos B cells were first transduced with the EGFP-CaaX retrovirus supernatant in a 1:1 ratio for 4 hr in a 37° incubator with 5%CO_2_. Cells were resuspended in fresh culture medium and incubated for another 7 days before sorting for the EGFP signal with a BioRad cell sorter. Sorted cells were transduced another round either with retrovirus encoding SNAP-IgM-BCR or SNAP-CD40 using the same protocol. Cells were stained with SNAP-Surface^®^ 549 (DY549P1) (S9112S, New England Biolabs) for 15 min in a 37° incubator with 5%CO_2_ and sorted for the fluorescent signal.

### Calcium flux measurements upon antigenic stimulation

1 million cells per mL were loaded with 1 μM Indo-1 (Catalog No. I1223, Invitrogen) in 0.1 % pluronic-containing cell culture medium for 30 min at a 37° incubator with 5% CO_2_. Loaded cells were washed in cell culture medium three times and rested in the incubator for 30 min. All calcium measurements with one batch of loading were completed within the 45 min following the 30 min resting period. Ratiometric measurements for Indo-1 were performed with BD Facs. The collected data points were saved as a .cvs file using the FlowJo export option. Calcium flux graphs were generated in Mathematica with a home-written code. Measurements were repeated at least 3 times and representative graphs were chosen for the manuscript.

### Sample preparation for LLSM and imaging

Lattice light-sheet microscopy was performed on a home-built clone of the original design by the Eric Betzig group [38]. Before imaging, 2 ml of a Ramos B cell suspension was centrifuged for 2 minutes at 700 x g. Afterwards, the supernatant was resuspended in 200 μl fresh cell culture medium together with 1 μM SNAP-Surface^®^ 549. After 15 minutes of incubation, the cell suspension was washed four times with 2 ml PBS, pH 7.4 at 37 °C. In the next step, the cell pellet was resuspended in approximately 20 μl PBS, pH 7.4 and mixed with phenol-red free Corning^®^ Matrigel^®^ Matrix (Product No. 356237, Corning) in a ratio of 1:1. 2 μl of this mix was dropped on a freshly plasma cleaned 5 mm round glass coverslips (Art. No. 11888372, Thermo Scientific). Coverslips were then transferred to a small humidified chamber and incubated at 37 °C until gel formation was complete. Then, the coverslip was mounted in a custom-built sample holder that was attached on top of a sample piezo. This ensures that the sample is inserted at the correct position between the excitation and detection objectives inside the sample bath containing PBS, pH 7.4 at 37 °C. For the inhibition of the Arp2/3 complex, CK-666 (Cat. No. 3950, Tocris) was added at a concentration of 100 μM to the sample bath. For the destabilization of the whole actin cytoskeleton, 2 μM Latrunculin A (Cat. No. 3973, Tocris) was added to the media bath. In both cases, cells were imaged for up to one hour. For the stimulation of cells via IgM-BCR, 50 ng/ml NIP-15-BSA (Biosearch Technologies) was added to the media bath and images were collected after a short incubation of about 5 min and proceeded for about 30 min. To test the antigen stimulation of cells in CK-666- or Latrunculin A-treated cells, cells were first incubated for 30 min inside the microscopy chamber with the respective substance before the addition of 50 ng/ml NIP-15-BSA. Imaging was then performed after 5 min and proceeded until 30 min. For image acquisition, a dual-channel image stack was acquired in sample scan mode by scanning the sample through a fixed light sheet with a step size of 500 nm which is equivalent to a ~ 270 nm slicing with respect to the z-axis considering the sample scan angle of 32.8°. A dithered square lattice pattern generated by multiple Bessel beams using an inner and outer numerical aperture of the excitation objective of 0.48 and 0.55, respectively, was used during the experiments. Channels were sequentially excited using a 488-nm laser (2RU-VFL-P-300-488-B1R; MPB Communications Inc., Pointe-Claire, Canada) for GFP and a 561 nm laser (2RU-VFL-P-2000-561-B1R; MPB Communications Inc.) for SNAP-Surface^®^ 549. Fluorescence was collected by a water dipping objective (CFI Apo LWD 25XW, NA 1.1, Nikon) and finally imaged on a sCMOS camera (ORCA-Fusion, Hamamatsu, Japan) with a final pixel size of 103.5 nm. Acquisition was performed at 100 frames per second with an exposure time of 8 ms for each channel.

### LLSM data processing

Lattice light sheet raw data were further processed by using an open-source LLSM postprocessing utility called LLSpy (https://github.com/tlambert03/LLSpy) for deskewing, deconvolution, channel registration and transformation. Deconvolution was performed by using experimental point spread functions recorded from 100 nm sized FluoSpheres™ (Art. No. F8801 and F8803, ThermoFisher Scientific) and is based on the Richardson–Lucy algorithm using 10 iterations. For channel registration and alignment, 200 nm sized fluorescent TetraSpeck™ microspheres (Art. No. T7280, ThermoFisher Scientific) were imaged with the same step size of 500 nm as used for cellular imaging. Alignment was done with a 2-step transformation method that deploys an affine transformation in xy and a rigid transformation in z.

### Image Analysis

#### Section 1: Segmentation of plasma membrane features

Binary masks for different topographical features of B-cell surface were created by using the following protocol:

- Creation of a binary mask for the entire plasma membrane surface using the GFP-CaaX signal.
- Creation of a binary mask for the inner space of the cell corresponding to the cytoplasm (named “dome” throughout the text).
- Refinement of the plasma membrane mask from step 1 using the dome from step 2.
- Creation of a binary mask for the ridge network using the GFP-CaaX signal.
- Partitioning of the plasma membrane topography into multiple morphological features using the binary masks from step 1, 2, 3 and 4.

##### Step 1: Binary segmentation of the plasma membrane surface

To perform the binarization of the GFP-CaaX signal, a 3D segmentation model by using the network architecture DynUNet was trained, which is implemented in Project MONAI (https://github.com/Project-MONAI/MONAI/tree/0.8.1). DynUNet itself is modified from nnU-Net developed by Isensee *et al.* [42]. To train this model, we first generated the ground truth dataset by using a semi-automatic segmentation approach. This approach consists of i) training an unsupervised binary voxel classification model that uses k-means clustering on multivariate feature space, and ii) manual correction of images segmented by this model and validation of the final binary masks by three domain experts. The ground truth dataset was generated from 6 x 3D images each extracted from a different LLSM time series. The feature extraction for training of the voxel classifier includes multiscale filtering of the image with a series of filters (filter names and filter parameters provided in **Table 1**). Manual correction of the preliminary binary masks produced by the voxel classifier was performed using the Napari package version 0.4.12 [59] as GUI. Prior to DynUNet model training, the following data augmentations were performed: cropping, contrast adjustments, Gaussian noise injections, Gaussian smoothing, Gibbs noise injections and rotations. The augmentations were performed by using the “transforms’’ module of the MONAI package. The DynUNet model was then trained based on the parameters and specifications provided in **Table 2**. The trained model was used to segment each volume in the volumetric time-course data. The output from this segmentation was refined by removing binary objects smaller than 30 voxels.

**Table 1:** Specifications of the filters used in the voxel-wise feature extraction.

| Filter Type | Parameter(s) | Value(s) |
| --- | --- | --- |
| Minimum | Window (Square Box) Dimension | 3 |
| Gaussian | Sigmas | 1, 1.5 |
| LoG | Sigmas | 0.8, 0.82, 0.84, 0.9, 0.93, 0.96, 1.0, 1.5 |
| DoG | Sigma1 and Sigma 2 | 5 and 7 |
| Grey Closing | Window (Sphere) Diameter | 5 |
| White Top-hat | Window (Sphere) Diameter | 3 |

**Table 2:** Specifications of the segmentation model. Any specifications not disclosed in this table are equal to the defaults implemented in the MONAI version 0.7.0 (Version 0.7.0 10.5281/zenodo.5525502)

|  |  |
| --- | --- |
| **Model Architecture** | DynUNet (https://docs.monai.io/en/stable/networks.html#dynamic-unet-block) |
| number of spatial dimensions | 3 |
| number of input channels | 1 |
| number of output channels | 2 |
| convolution kernel size | (1, 3, 3, 3) |
| convolution strides<br>(=upsample kernel size) | (1, 2, 2, 2) |
| dropout | None |
| normalisation | instance normalisation |
| activation layer | leakyrelu |
| deep supervision | False |
| residual connections | True |
| **Loss function** | DiceCELoss (https://docs.monai.io/en/stable/losses.html#diceceloss) |
| to_onehot_y | True |
| softmax | True |
| squared_pred | True |
| batch | True |
| **Optimiser** | Novograd (https://docs.monai.io/en/stable/optimizers.html#novograd) |
| learning rate | 1e-2 |

**Table 4:**
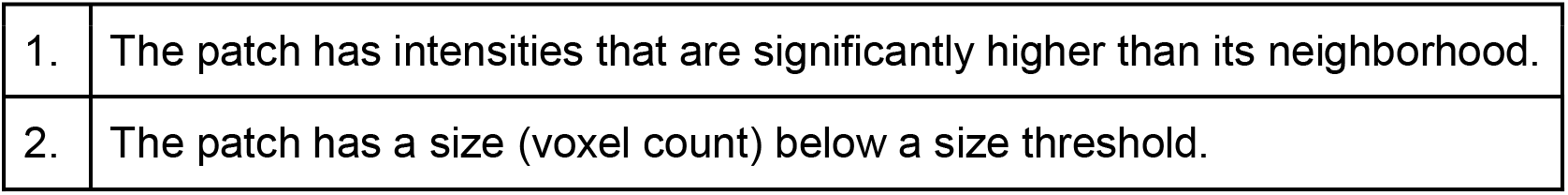
Cluster criteria:

|  |  |
| --- | --- |
| 1. | The patch has intensities that are significantly higher than its neighborhood. |
| 2. | The patch has a size (voxel count) below a size threshold. |

##### Step 2: Extraction of the dome

A semi-automatic, robust, watershed-based protocol was developed and applied to extract the dome of the cell directly from the GFP-CaaX signal. The protocol is based on automatic selection of two seeds for the interior and exterior of the cell. The algorithm is outlined as follows:

i. Binarize the raw image with a low threshold to obtain a rough binary surface object.
ii. If the surface object contains gaps, use mathematical morphology to close them. Then fill the interior space of the object via a binary filling.
iii. Use binary opening to eliminate any protrusive structures from the filled surface. This leads to a rough dome-like structure. Extract the surface layer of this dome using either mathematical morphology or specifying a level of its distance transform.
iv. Compute a distance transform using the surface layer from step iii as the zero level.
v. Select a distance threshold to specify internal and external seeds from the distance map. For example, a distance threshold of 3 μm will specify an internal patch 3 μm inwards from the surface and an external shell enclosing the cell from 3 μm outwards from the surface. Label the internal and external seeds.
vi. Generate an input image for the watershed algorithm by calculating a weighted mean of the raw image and the distance map from step iv. The weights can be specified by the user.
vii. Perform the watershed segmentation on the input image using internal and external labels as markers. The dome is the component corresponding to the internal label in the output.

##### Step 3: Refinement of the plasma membrane mask

Plasma membrane mask from (**section-1 / step-1**) usually contains small gaps at the cell surface, which may interfere with the downstream analysis. To close these gaps, we merged the output from **(section-1 / step-1)** and the surface layer of the dome from (**section-1 / step-2**). We subtracted the entire dome object except its surface layer, from the plasma membrane mask, in order to ensure that the downstream analysis is confined to the cell surface and the microvilli, and that any other components within the cytoplasm are excluded from the analysis.

##### Step 4: Extraction of the ridge mask and the skeleton

Hessian-based approaches are frequently used for detection and segmentation of curvilinear objects [60]. Here the following pipeline was used for extraction of the ridges:

i. Apply a Hessian-based vesselness filter [61] (implemented in ITK version 5.2.1) to the intensity image.
ii. Rescale the values of the Hessian response image between 0 and 1.
iii. Binarize the rescaled Hessian response image using a combination of local and global thresholding, where the local threshold *t* is defined as:

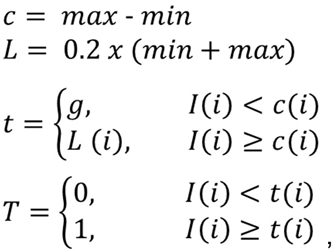

where,

*I* is the image array and *i* is the voxel index.
*max* and *min* are the local maxima and local minima transforms of the image, respectively, calculated with a spherical local window having a diameter of 7 voxels;
*g* is a user-specified global threshold. Here we used multi-otsu thresholding [62], implemented in scikit-image version 0.18.3) to calculate 4 thresholds and assigned the lowest threshold to *g*;
*T* is the final thresholded image.
iv. Subject the thresholded image to one round of closing operation with a spherical structuring element in a window of (3, 3, 3). The output from this process is the ridge mask.
v. Subject the binary ridge mask from step iv recursively to the steps i to iv. This yields a thinned form of the ridge mask.
vi. Apply 3D morphological skeletonization [63], implemented in scikit-image version 0.18.3) to the thinned ridge mask from step v. The output from this process is the 1-voxel thick ridge skeleton.

##### Step 5: Partitioning of the plasma membrane mask into its topographical features

###### Step 5A: Partitioning of the cells that are not treated with latrunculin

By using the outputs from (**section-1 / steps-2, 3 and 4**), the topographical features of the plasma membrane mask were identified. This process is further divided into multiple steps as described below.

**Step 5A/a**: Separating the protrusive structures (microvilli) from the cell body:

The pipeline, which is based on binary mathematical morphology, is outlined below:

i. Calculate the union of the dome from **section-1 / step-2** and the plasma membrane mask from **section-1** / **step-3** to obtain an “in-filled” cell mask.
ii. Apply a binary opening filter to the output from step i using a box-shaped structuring element. We used a structuring element of voxel size (3, 5, 5).
iii. Apply a binary opening filter to the output from step ii using an ellipsoidal structuring element. We used a structuring element of voxel size (5, 7, 7).
iv. Calculate the intersection of the output from step iii and the plasma membrane mask. This process yields the cell body mask.
v. Subtract the cell body mask from the plasma membrane mask. This process yields the microvilli mask.

**Step 5A/b**: Extraction and annotation of the ridge network:

A custom algorithm was applied to the ridge skeleton from **section-1 / step-4** to detect and label microvilli tips and ridge junctions. In the first step of the algorithm, the skeleton vertices are annotated based on the local sum in a (3, 3, 3) neighborhood of each vertex as described in the following:

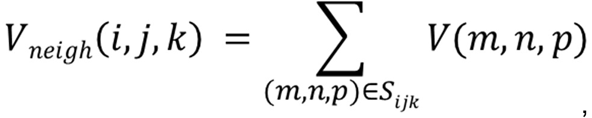

where,

*V* is the skeleton image,

*V_neigh_* is the local sum evaluated over the skeleton,

*m, n* and *p* are the coordinates in the local window of (3, 3, 3) around the point (i, j, k) *i, j* and *k* are the global coordinates

The annotation is then performed as described below:

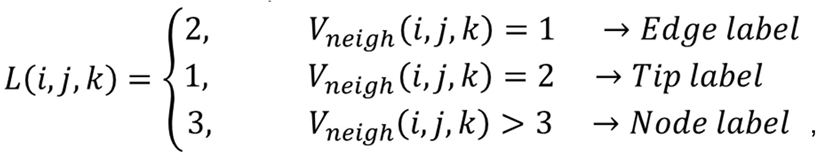

where,

*L* is the label image.

In the second step of the algorithm, a correction function to the label image was applied, which recognizes very short edges (<200nm) and replaces them with a node label. We then used our feature masks from **section-1 / step-5A/a** to confine the annotations. Namely, the cell body mask was used to confine the nodes and edges to the cell body and the microvilli mask to confine the tips to the microvilli.

**Step 5A/c**: Construction of the feature image:

Three geodesic distance maps per cell were computed based on the labeled network image from **section-1 / step-5A/b** using the fast marching method (implemented in scikit-fmm version 2021.10.29). The first geodesic map (gdmap_skeleton) was computed by assuming the intersection of the skeleton and the cell body mask as the zero level set and the cell body mask as the speed image. The second geodesic map (gdmap_tips) was computed by assuming the tips as the zero level set and the microvilli mask as the speed image. Finally, the third geodesic map (gdmap_nodes) was computed by assuming the nodes as the zero level set and the cell body mask as the speed image.

After computing the geodesic distance maps, the final topographical features were computed as follows:

- Microvilli tips mask was obtained by applying a threshold of 2.1 to the gdmap_tips.
- Microvilli shafts mask was obtained by subtracting the microvilli tips mask from the microvilli mask.
- Cell body node mask was obtained by applying a threshold of 1.5 to the gdmap_nodes.
- Cell body ridge mask was obtained by applying a threshold of 2.7 to the gdmap_skeleton and then subtracting the cell body node mask from the resulting binary mask.
- Cell body shallow invaginations mask was obtained by subtracting the union of cell body ridge mask and the cell body node mask from the cell body mask

###### Step 5B: Partitioning of the latrunculin-treated cells

Cells that were treated with latrunculin displayed a generally smooth surface with occasional, bulky stacks of membrane. As these cells did not show an organized ridge network, they were not suitable for partitioning using the approach described in **section-1 / step-5A**. The latrunculin-treated cells showed a surface topography that contained short stretches of curvilinear elevations as well as disorganized blob-like chunks of membrane with varying sizes. To detect both types of structures, two detection filters to each raw image were separately applied, one for curvilinear structures [64] and one blob-shaped structures [65]. To detect structures of different sizes, each filter was applied at multiple scales, using sigma values of 1 and 2 for the curvilinearity filter and sigma values of 1, 2 and 3 for the blob filter. Then each response image was segmented using a global threshold, which was generally chosen to be the Otsu threshold [66] but varied when necessary. When the binary masks were obtained for both curvilinear and blob-shaped structures, these were combined using a Boolean union filter to obtain a preliminary “elevation” mask. Finally, a watershed segmentation was applied to delineate the elevation mask. For this watershed step, we selected the preliminary elevation mask as the marker 1, the dome from **section-1 / step-2** as the marker 2 and the negative of the Euclidean distance transform of the cell surface mask from **section-1 / step-3** as the main input image. As a result of the Watershed segmentation, the label corresponding to the marker 1 was designated as the elevation mask and the label corresponding to the marker 2 was designated as the cell body mask.

#### Section 2: Segmentation of IgM Clusters

It was asserted that a voxel patch must meet the following criteria to be assumed as a cluster:

In order to extract patches that meet criterion 1, a two-step algorithm was developed, which uses a custom local thresholding function with hyperparameters adaptive to different images. The two steps are outlined as follows:

##### Step 1: Extract a “blob” mask

A blob mask is an image, which is binarized based on local curvature values calculated from the raw image. The local curvature values are calculated based on the eigenvalues of the Hessian matrix *H(f)*:

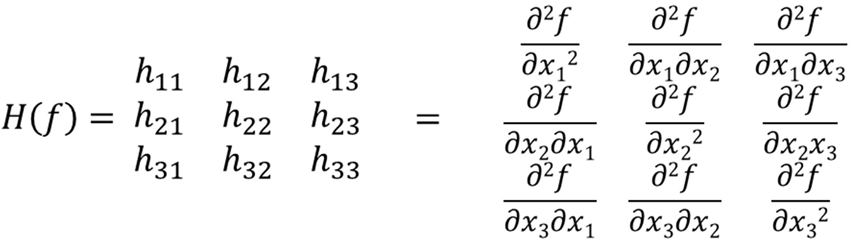

where *f* is convolution of the image *I* with a Gaussian kernel (we used a sigma value of 1). The eigenvalues *λ*_1_, *λ*_2_ and *λ*_3_ were calculated from the *H(f)*. Subsequently, the blob mask *B* was calculated as below:

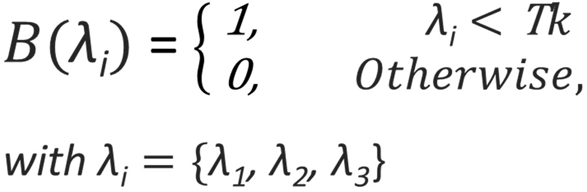

Where T is the Otsu threshold of the <u>negative</u> values of λ**i** and *k* is a user-specified value (we selected the value of 0.01 for an over-segmentation). Finally, the blob mask B was calculated as the intersection of the components:

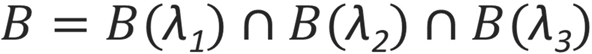

The blob mask calculated in this manner contains nearly all possible voxel patches that can be extracted from a grayscale image *I*. These patches are treated as “candidates” to be clusters.

##### Step 2: Find optimal local thresholding hyperparameters

A local thresholding function based on a modification of the Niblack threshold method [67] was used in that the binarization via local thresholding is restricted to the high-contrast regions in the image as expressed below:

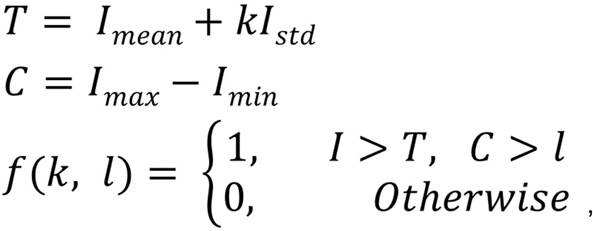

where *I_mean_*, *I_std_*, *I_max_*, *I_min_* are arrays with local means, standard deviations, maxima and minima, respectively, calculated using a square-box local neighborhood with window size of 5 voxels. *T* is the Niblack threshold and *C* is the local contrast. The segmentation function *f* is then dependent on the hyper-parameters *k* and *l,* which must be optimized for segmenting the clusters of a particular image. To this end, we applied a grid search for a series of *k* and *l* and chose the binary mask with greatest similarity to the blob mask *B* from (**section-2 / step-1**). In other words, for a specific image I, we applied:

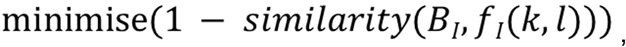

where we used the Tversky method [68] as the similarity function between the two binary images.

##### Step 3: Apply a size threshold

The output image from (**section-2 / step-2**) was filtered with a size threshold so that binary objects with a voxel count greater than a user-specified threshold are ruled out from being considered clusters. A size threshold of 75 voxels was selected for this step and the output was designated *M_c_* (cluster mask).

##### Step 4: Locate centroids and apply feature-based contrast threshold

The centroid of each cluster from (**section-2 / step-3**) was located and labeled and designated *C_c_* (cluster centroids). Subsequently, the ratio of the IgM intensity at each cluster centroid to the mean intensity of the corresponding morphological feature was calculated. The output is a map of feature-based intensity fold increase at cluster centroids. Finally, a threshold was applied to this map so that any centroids with a ratio lower than a given threshold are deleted. Depending on the analysis, this threshold was specified as 1.25 or 3, meaning that each centroid must have an intensity 1.25 fold (for moderate stringency) or 3 fold (for high stringency) higher than the mean intensity of the corresponding morphological feature.

### Analysis

#### Granularity

Granularity was defined as the ratio of the voxel count of the size-filtered binary objects (**section-2 / step-3**) to the total voxel count of the binary objects yielded by the cluster segmentation protocol (**section-2 / step-2**).

#### Feature-based cluster density

Feature-based cluster density is the ratio of the cluster count within a particular morphological feature to the total volume of that feature.

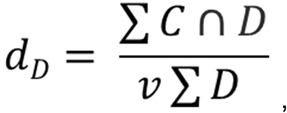

where, *d_D_* is the cluster density for a particular morphological feature. *D* is the mask for the morphological feature in question. *C* is the centroid mask and *v* is the voxel volume for the image.

#### Cluster density plotted with geodesic distance

To analyze the cluster density in relation to geodesic distance, the geodesic distance maps from (**section-1 / step-5A/c**) were divided into equal-sized bins so that each bin represents a particular geodesic distance range. Then the cluster density was calculated within each individual bin as described below:

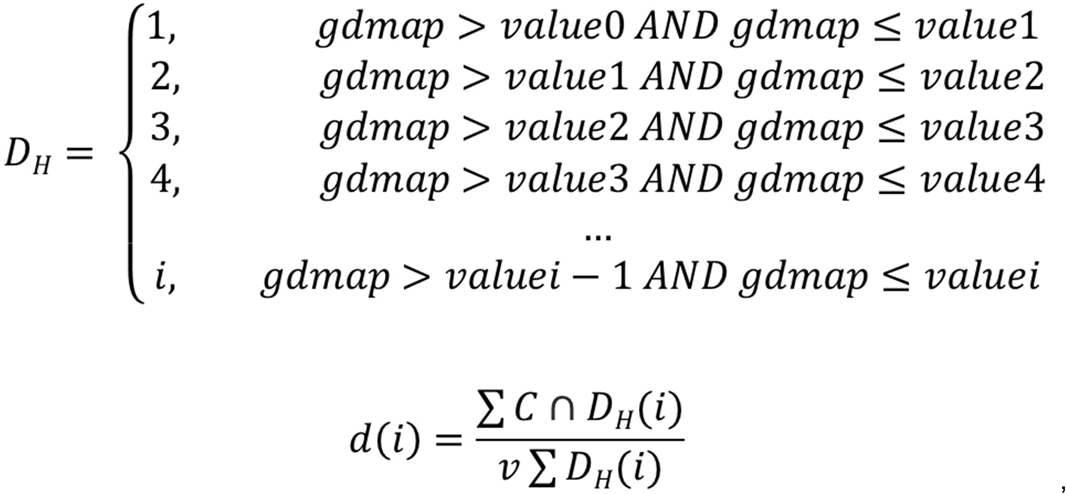

where *D_H_* is the binned image, *i* is the index to a particular bin (hence represents geodesic distance), *d(i)* is the density calculated for the particular bin indexed by *i*.

#### Global cluster enrichment

Global cluster enrichment is defined as the fold increase of global mean value of clustered IgM signal over global mean value of non-clustered IgM signal:

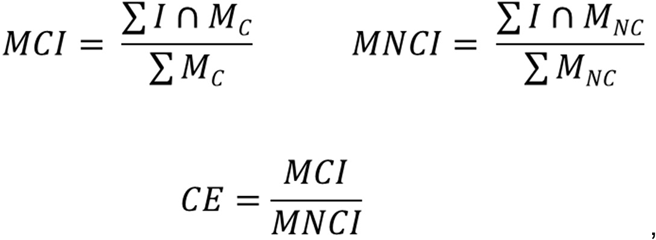

where *I*: grayscale image

*M_C_*: cluster mask

*M_NC_*: non-clustered surface mask

MCI: mean clustered intensity

*MNCI*: mean non-clustered intensity

*CE*: cluster enrichment

#### Cluster enrichment per feature

Cluster enrichment per feature is defined as the fold increase of mean value of clustered IgM signal within a particular morphological feature over global mean value of non-clustered IgM signal:

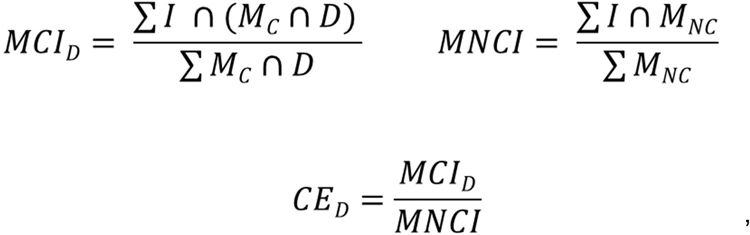

where *D*: binary mask for a specific morphological feature

*MCI_D_*: mean clustered intensity within feature

*CE_D_*: cluster enrichment for feature *D*

#### Proportion of the clustered intensities

To calculate the proportion of clustered signal, the summed value within the cluster mask was divided to the summed value within the entire surface mask (clustered + non-clustered):

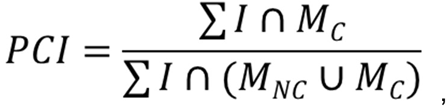

where *PCI*: proportion of the clustered intensities

#### Calculation of staticity for cluster centroids

To create a staticity map, a 4D label image (the volumetric time-course) for the cluster centroids was correlated with a 4D Gaussian kernel. This correlation operation yields a stronger response with cluster centroids that remain within a close neighborhood compared to centroids that undergo large displacements. In other words, voxels corresponding to relatively immobile centroids have high values in the map, whereas voxels corresponding to highly dynamic centroids have low values. Once we had the staticity map, we calculated the staticity as the proportion of voxels higher than a user-defined threshold. Therefore:

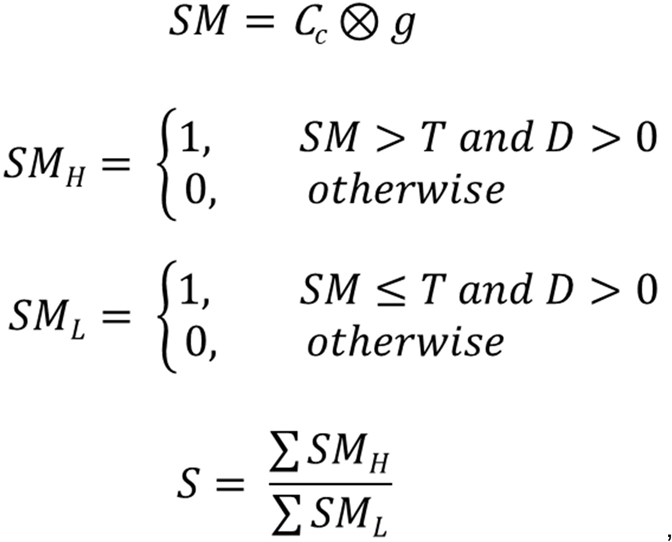

where *C_c_*: the binary input image for cluster centroids, which is subjected to the staticity analysis,

*SM*: staticity map,

*g*: Gaussian kernel with shape (7, 5, 5, 5) voxels and weights (3, 1, 1, 1) in the order of (t, z, y, x),

*S*: the staticity term

The staticity was calculated for the first 15 frames of each time-course image.

#### Calculation of staticity for surface features

With label images of surface features, staticity was calculated based on the similarity of consecutive time frames. To measure similarity, the intersection over union (IOU) method was used. Highly dynamic surface features undergo large displacements between time frames resulting in low IOU scores, whereas relatively immobile features yield high IOU scores. The staticity is then the mean of the IOU scores calculated for all consecutive for all consecutive pairs along the first 15 frames of each time-course dataset.

Therefore:

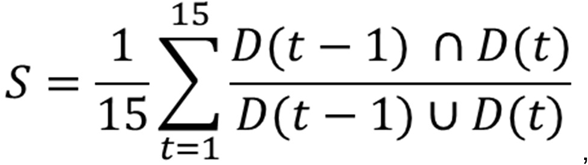

where *S*: the staticity term,

*D*: the binary input image for one of the computed surface features, which is subjected to the staticity analysis,

*t*: the time frame

#### Statistics and Reproducibility

Statistical analyses were performed using GraphPad Prism. Experiments were performed independently, repeated at least twice and similar results were obtained. Data was only excluded when the image quality did not suffice for image analysis. Distribution was assumed as normal for statistical analysis. Data collection was not performed blind. For statistical analyses, either two-tailed paired or unpaired Student’s ŕ-tests were performed, as specified in the figure legends. Significance was drawn at * p < 0.05. Significance was defined as **** p < 0.0001, *** p < 0.001, ** p < 0.01.

## Author Contributions

D.S. and M.R. designed the research. D.S. performed the experiments. J.P. and R.K. implemented the LLSM platform and M.H. performed the LLSM imaging and data postprocessing. B.Ö. developed the image analysis tools and analyzed the LLSM data. R.R., R.K., J.P. provided collaborative support. D. S. and M.R. wrote the manuscript with input from all coauthors.

