## Supplemental Information for "Plasma membrane topography governs the three-dimensional dynamic localization of IgM B cell receptor clusters"

**Supplemental Figure 1:**

**Related to Figure 1**

**(a)** Model of the NIP-specific SNAP-tagged mIgM molecule expressed on H/L-deficient Ramos B cells via a retroviral expression vector carrying the cDNA sequence of a SNAP-tagged μ H-chain separated by a P2A element from the sequence encoding the λ L-Chain. The N-terminal SNAP-tag is preceded by a leader sequence (LS) and separated from the start of the $V_H$ sequence by a 17 amino acid long linker with the sequence $A(EAAAK)_3A$. To visualize the receptors specifically at the B cell surface, the membrane-impermeable benyzlguanine-DY549P1 was covalently attached to the SNAP-tag. **(b)** Cross-section widefield images of EGFP-CaaX- and EGFP-CaaX/SNAP-IgM-BCR-transduced H/L-KO Ramos B cells at the midline. SNAP-IgM-BCR is visualized with the BG-DY594P1 SNAP-tag substrate. **(c)** Calcium influx response of SNAP-IgM-BCR-expressing H/L-KO Ramos B cells loaded with Indo1. Cells are stimulated with the indicated molecules. **(d)** Cross-section widefield images of SNAP-IgM-BCR-expressing H/L-KO Ramos B cells stimulated with Streptavidin-649 pre-incubated NIP-15-BSA-biotin (50 ng/ml) followed by PBS wash**. (e)** Overview of the segmentation protocol for the whole cell surface (introduced in fig. 1) and cell surface features (introduced in fig. 2 and 3). Steps are numbered as explained in the methods section.

**Supplemental Figure 2:**

**Related to Figure 3**

**(a)** Volume distribution of the different surface features as computed from the surface mask of untreated cells. **(b, c, d)** Fold change in the mean intensity of EGFP-CaaX **(b)**, IgM-BCR **(c)**, and CD40 **(d)** within the R+N and MV feature masks normalized to the mean intensity at the SI feature mask. **(e, f, g)** Continuous analysis of CD40 cluster density using geodesic distance maps initiated from **(e)** MT towards MV roots, **(f)** centerline of R towards cell body surface, and **(g)** N towards cell body surface. n = 7 for EGFP-CaaX and IgM-BCR. n =3 for CD40.

**Supplemental Figure 3:**

**Related to Figure 4**

**(a)** Phalloidin-Alexa555 staining of fixed and permeabilized Ramos B cells expressing EGFP-CaaX and IgM-BCR with or without 30 min 100 μM CK-666 treatment. **(b)** Node degree analysis of the skeletonized R network in untreated and 100 μM CK-666-treated cells for up to 30 min. n = 7 for untreated and n = 4 for CK-666. **(c)** Sum projections of N and skeletonized R network (excluding the MV) in an untreated and 100 μM CK-666-treated cell for 15 consecutive time frames (duration: 37.5 s).

**Supplemental Tables:**

Supplemental Table 1: Microvilli-related genes expressed in Ramos B cells

Supplemental Table 2: Microvilli-related genes not expressed in Ramos B cells

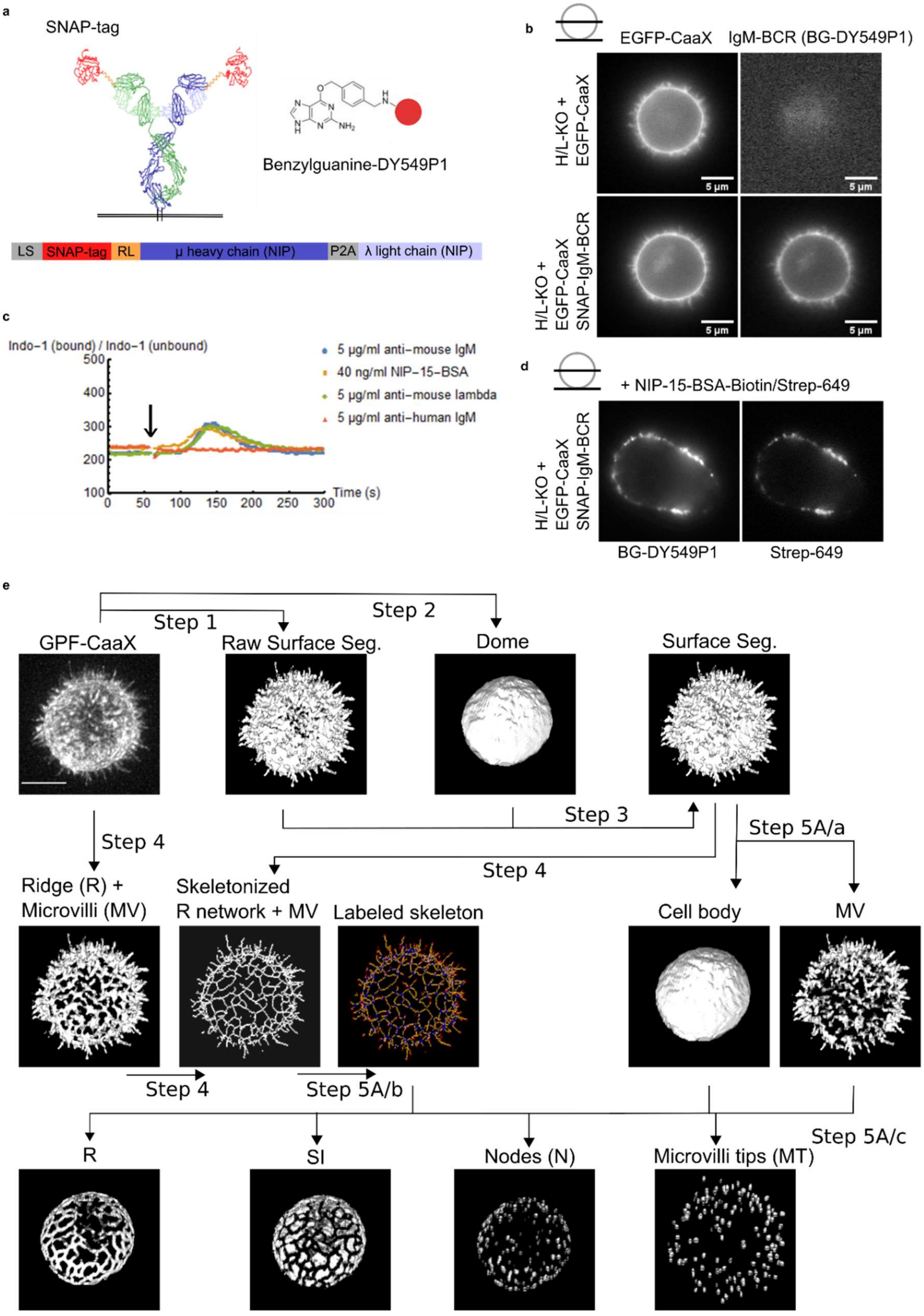


**Supplemental Figure 1**

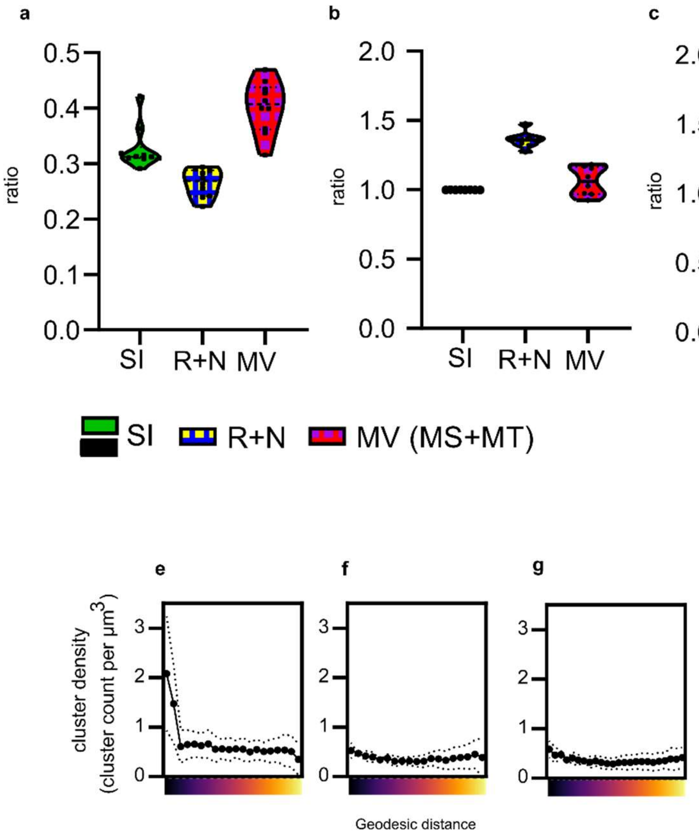


**Supplemental Figure 2**

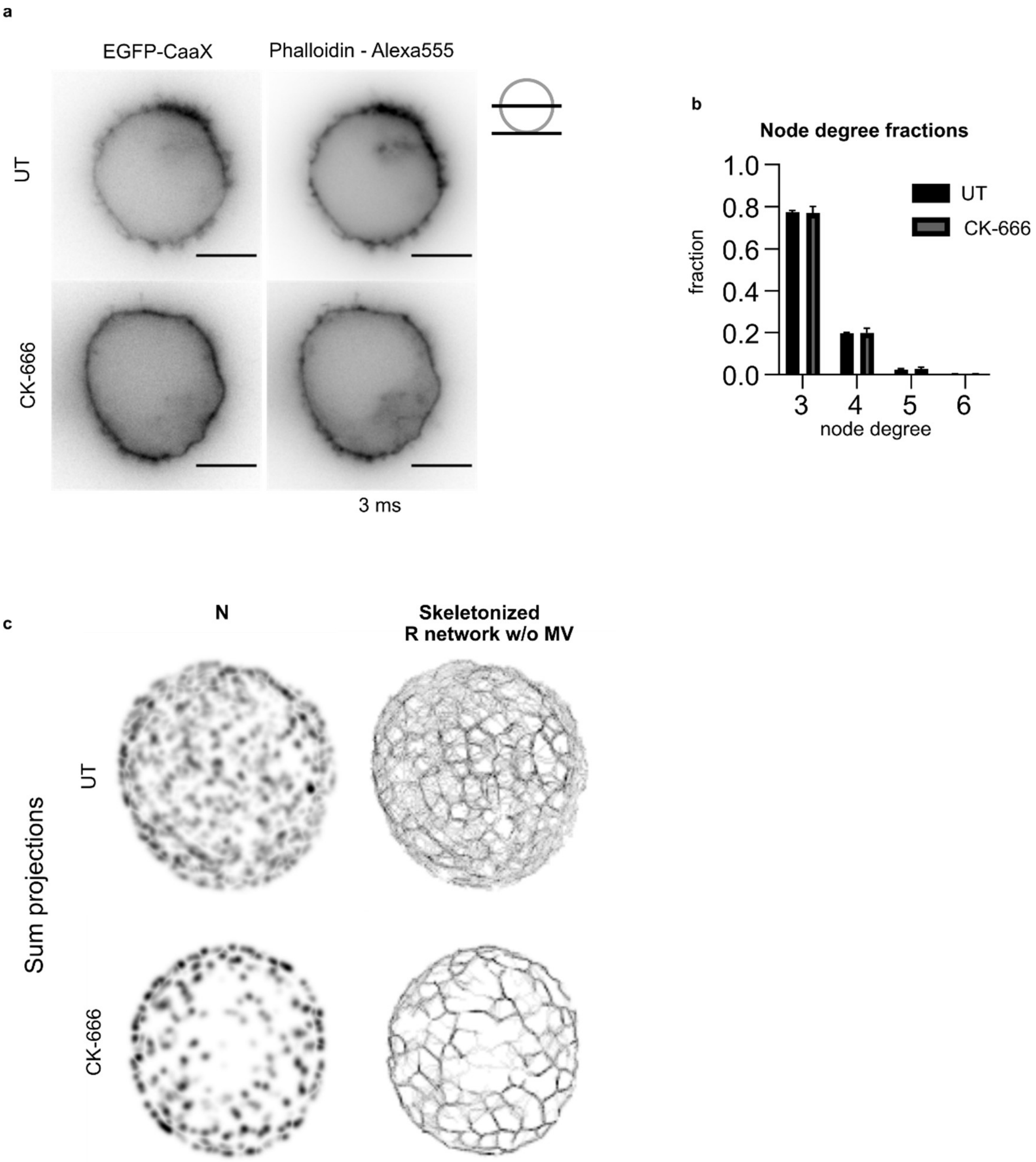


**Supplemental Figure 3**

**Supplemental Table 1:**
Microvilli-related genes expressed in Ramos B cells:

| Gene | Raw Transcript | Reference |
|---|---|---|
| ACTR2 | 1359 | [1] |
| ACTR3 | 904 | [1] |
| ARF6 | 509 | [1] |
| ARPC1B | 853 | [1] |
| ARPC4 | 516 | [1] |
| ASAP1 | 33 | [2] |
| BSG | 1075 | [1] |
| CDC42 | 892 | [1] |
| CFL1 | 5631 | [1,3] |
| CLIC1 | 1721 | [1] |
| CLIC4 | 341 | [1] |
| CD81 | 2494 | [1] |
| DSTN | 1222 | [1] |
| EPS8L1 | 20 | [4] |
| EZR | 2424 | [1,5,6] |
| GNAI2 | 798 | [1] |
| GNAI3 | 393 | [1] |
| GNB1 | 1996 | [1] |
| GNB2 | 1801 | [1] |
| GNB4 | 281 | [1] |
| ITGAL | 459 | [1] |
| MSN | 394 | [1,7] |
| MYO1D | 92 | [8] |
| MYO1G | 524 | [1,9] |
| MYO3B | 238 | [10] |
| MYO6 | 282 | [11] |
| PFN1 | 6069 | [12] |
| PLS1 | 15 | [13] |
| PLXNA1 | 236 | [1] |
| RAB8B | 133 | [1] |
| RAB35 | 248 | [1] |
| RAP1A | 2079 | [1] |
| RAP1B | 502 | [1] |
| RDX | 43 | [1] |
| RHOA | 1659 | [1] |
| RRAS2 | 130 | [1] |
| SLC16A1 | 321 | [1] |
| SLC3A2 | 1453 | [1] |
| SLC9A3R1 | 549 | [1,14] |
| SLC7A5 | 1275 | [1] |
| TAGLN2 | 1994 | [15] |
| TRPV2 | 95 | [1] |
| WHRN | 193 | [16] |

**Supplemental Table 2:**
Microvilli-related genes not expressed in Ramos B cells:

| Gene | Raw Transcript | Reference |
|---|---|---|
| BAIAP2 | 9 | [17] |
| BAIAP2L1 | 1 | [18] |
| CD5 | 0 | [1] |
| CD9 | 1 | [19] |
| CD44 | 1 | [1] |
| CLIC5 | 0 | [11] |
| COBL | 1 | [2] |
| ENPEP | 3 | [1] |
| EPS8 | 1 | [20] |
| ESPN | 1 | [21,22] |
| GNAI1 | 0 | [1] |
| FSCN1 | 0 | [23] |
| MYO1A | 1 | [24] |
| MYO3A | 0 | [25] |
| MYO5B | 0 | [26] |
| MYO7A | 0 | [27] |
| MYO7B | 4 | [28] |
| MYO15A | 2 | [16] |
| PROM1 | 1 | [29] |
| VIL1 | 1 | [30] |
